# A cortical immune network map identifies a subset of human microglia involved in Tau pathology

**DOI:** 10.1101/234351

**Authors:** Ellis Patrick, Marta Olah, Mariko Taga, Hans-Ulrich Klein, Jishu Xu, Charles C White, Daniel Felsky, Chris Gaiteri, Lori B Chibnik, Sara Mostafavi, Julie A Schneider, David A Bennett, Elizabeth M Bradshaw, Philip L De Jager

## Abstract

Microglial dysfunction has been proposed as one of the many cellular mechanisms that can contribute to the development of Alzheimer's disease (AD). Here, using a transcriptional network map of the human frontal cortex, we identify five gene modules of co-expressed genes related to microglia and assess their role in the neuropathologic features of AD in 541 subjects from two cohort studies of brain aging. Two of these transcriptional programs – modules 113 and 114 – relate to the accumulation of β-amyloid, while module 5 relates to tau pathology. These modules are also detectable in the human brain's epigenome, where we replicate these associations. In terms of tau, we propose that module 5, a marker of activated microglia, may lead to tau accumulation and subsequent cognitive decline. We validate our model further by showing that *VASP*, a representative module 5 gene, encodes a protein that is upregulated in activated microglia in AD.

## Introduction

Alzheimer’s disease (AD) is characterized pathologically by the accumulation of both β-amyloid and tau pathologies which lead to the gradual loss of cognitive function and, ultimately, dementia^1^. The amount of these two pathologies that is present in the older brain is strongly but only partially correlated^2^, enabling us to distinguish molecular pathways that are involved in one or the other process. While genome-wide association studies have unequivocally pointed to the innate immune system and particularly myeloid cells as major contributors to AD pathophysiology ^3-5^, our current understanding of the mechanistic involvement of microglia and infiltrating macrophage in human AD pathology is rudimentary as it is largely based on studies performed in animal models of AD, which imperfectly capture aspects of human disease. Myeloid cells have been implicated in all aspects of AD, from the asymptomatic phase of amyloid accumulation to the progression of dementia, but there is little clarity on how disparate human observations can be assembled into a coherent picture. Therefore, to identify the different components of human myeloid responses that exist in the neocortex and to examine how each component contributes to the continuum of AD-related pathological and clinical traits in the aging brain, we evaluated a recently derived Molecular Network Map of the aging human frontal cortex using tissue level RNA sequence (RNAseq) data obtained from the frozen dorsolateral prefrontal cortex of participants in two prospective studies of cognitive aging, the Religious Order Study (ROS) and the Memory Aging Project (MAP)^6-8^. From the RNAseq data, we derive groups of co-expressed genes – that we term "modules" – and build the network from these modules as well as outcome measures available from each of the 541 participants that were profiled. As these individuals are non-demented at study entry, they represent a sampling of the older, aging population. At the time of death, they display a full spectrum of clinicopathologic features related to AD that are found in older people including cognitive decline, dementia, extracellular β-amyloid deposition, hyperphosphorylation of tau, and microglial activation^6,7,9^.

Here, using our recently established gene expression signature of aged human microglia^10^, we identify five microglia-related gene co-expression modules that capture different transcriptional programs of microglia. We focus on dissecting the role of these modules in AD; in mapping the conditional relationship between modules and cognitive and neuropathologic outcomes, we identify and validate a microglial module that contributes to the accumulation of tau pathology. Two other modules relate to β-amyloid, and a fourth – enriched for LOAD susceptibility genes – appears to be primarily related to aging. Thus, we provide an initial immune network perspective of the divergent sets of co-expressed genes that govern microglia identity in the aging human brain and identify the relationship of these components to specific pathologies.

## Results

### A molecular network map derived from RNA sequence data

ROS and MAP are two large longitudinal studies of aging that were designed and are managed by the same group of investigators so that their data can be merged in joint analyses ^11,12^. Participants are non-demented at study entry, agree to brain donation at the time of death, and are evaluated annually with a battery of 21 neuropsychologic tests. A person-specific slope of cognitive decline is calculated for each participant based on 17 tests in common^6,7^. Demographic and clinical characteristics of the 541 participants used in these analyses are presented in **Supplementary Table 1**.

To reduce the dimensionality of the RNAseq data generated from the DLPFC (Dorsolateral Prefrontal Cortex) of each subject, we previously defined modules of co-expressed genes using the Speakeasy algorithm ^13^ : there are 47 such modules that contain a minimum of 20 genes and a median of 331 genes. Thus, each module contains a group of genes that have a shared regulatory architecture, and the 47 modules represent the nodes of our network^8^.

### Identifying microglia-related modules

In this manuscript, we take a deeper look at the subset of these modules that capture the role of microglia. We identified modules enriched for microglial genes as defined in a new reference RNAseq data set derived from live, purified human microglia/macrophages extracted from fresh autopsy DLPFC samples of 10 ROSMAP participants of advanced age: the HuMi_Aged gene set consists of 1,081 microglia enriched genes that were identified based on a four-fold increase in expression in microglia vs. DLPFC expression^10^.

Of the 47 modules, five are enriched (p<0.0011) for the HuMi_Aged gene set: modules m5, m113, m114, m115, and m116 (**Table 1, Figure 1, Supplementary Figure 1**). Results are similar if we use other human microglial profiles^14^. To determine whether these enriched modules are specific to microglia, we performed a similar enrichment analysis using reference RNAseq profiles of human iPSC-derived neurons and primary human astrocytes (https://www.synapse.org/#!Synapse:syn2580853/wiki/409840). Of the five modules enriched for microglial genes, none are enriched for neuronal genes, but two modules (m113 and m115) are also enriched for astrocyte genes (**Supplementary Table 2**). The latter modules may relate to immune responses shared with astrocytes, which are well-known to contribute to central nervous system inflammatory responses^15^. By contrast, m5, m114 and m116 appear unique to microglia, with m116 displaying, by far, the greatest microglial enrichment: 67% of its 224 genes are present in our list of 1081 microglial genes. It contains many well-known myeloid/microglial genes such as *TREM2* and *TYROBP.* This result is emphasized in a parallel enrichment analysis of microglial, neuronal and astrocytic profiles generated from mouse brain ^16^ where we observe that m116 is the most cell-type specific of all tested modules (**Supplementary Figure 2**). Given the strength of its enrichment for genes that are reported to be microglial, m116 probably represents a set of genes expressed by all microglia. We note that m116 captures the same set of coexpressed genes that led to the report on the role of *TYROBP* in AD^4^.

**Figure 1.**
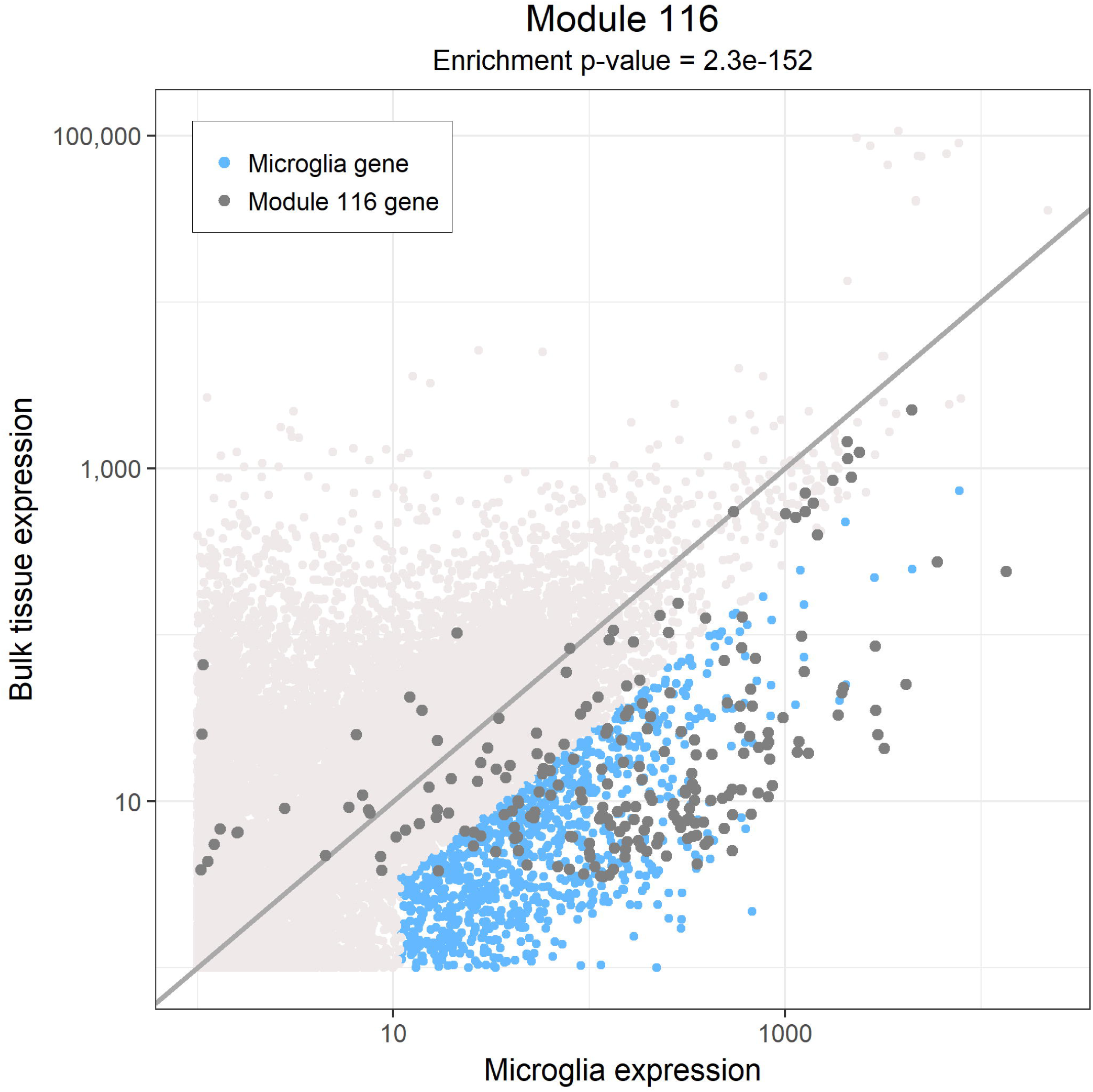
Identification of microglia related genes and modules. The expression (FPKM) of genes in bulk DLPFC tissue are compared to their expression in isolated microglia. We highlight microglia related genes in blue, these genes have a 4-fold higher expression in isolated microglia compared to bulk tissue are likely involved in processes that are relatively specific to microglia. The genes in module 116 are highlighted in gray and mostly appear to be more highly expressed in isolated microglia compared to bulk tissue.

**Table 1.**
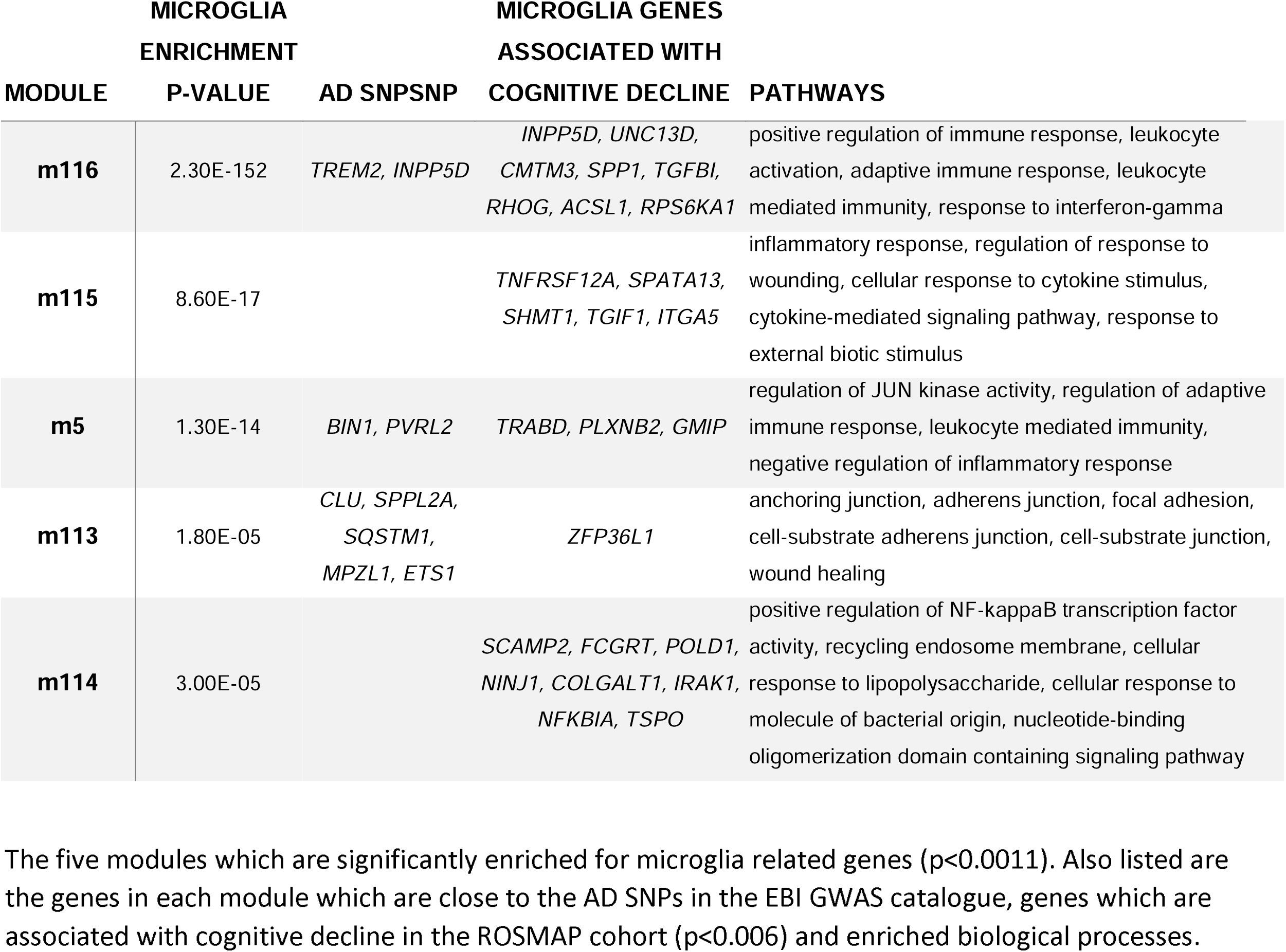
Enrichment of gene co-expression modules for microglia related genes

In our original evaluation of neocortical modules^8^, these five microglial modules were all robust: the component genes of each module (see **Supplementary Table 3**) are (1) co-expressed in an independent, publicly available frontal cortex RNA expression dataset^4^ and (2) correlated in H3K9Acetylation (a histone mark of actively transcribed genes) ChIP-seq data from an overlapping set of 669 ROSMAP individuals^8^. To obtain an initial perspective on their function, we annotated each module by identifying both known pathways enriched in each module (**Table 1, Supplementary Table 4**) and transcription factors whose binding sites are enriched in the promoters of each module’s genes (**Supplementary table 5**). For example, m5 is enriched for genes found in genesets related to c-JUN N-terminal (JNK) kinase activity, which is consistent with an enrichment of AP1 binding sites in the promoters of m5 genes. This enrichment appears to be specific to m5. Similarly, signatures of phagocytosis are found only in m116, while m113 appears to be specifically enriched for genesets involved in cell adherence and vascular development. Thus, m113 may reflect a transcriptional program active in perivascular astrocytes and myeloid cells since it is found in both cell types. By contrast, NFκB-related pathways (enriched in m114 and m115) and responses to type I interferons or interferon γ (enriched in m114 and m116) are split between different modules. The transcription factor binding site analysis is consistent with these results, with enrichment in interferon response factor (IRF) binding sites in m116 and in signal transducer and activator of transcription (STAT) binding sites in m115. Notably, the binding site for the SP1/PU.1 transcription factor implicated in AD susceptibility^17^ is enriched in m116 and also, more marginally, in m5. We observe that each module is enriched for unique subsets of transcription factors (**Supplementary Figure 3**) suggesting that each module may be maintained by an exclusive set of regulatory mechanisms.

We previously reported that m116 is enriched in AD susceptibility genes^18^ (**Supplementary Table 6**), including *TREM2* and *INPP5D* (**Table 1**)^19^. While not enriched over the background, other modules also contain genes found in AD loci, such as *BIN1* in m5. These genetic associations emphasize the strong causal role of myeloid and microglial cells in AD pathogenesis.

### Modules diverge in their association with AD pathologies

To resolve the relative roles of the different modules in different aspects of AD, we determined their relation to the rich clinicoalpathologic phenotypes available in ROSMAP participants using a meta-feature calculated for each of the five modules. In **Table 2**, we find that m5 and m114 display the most associations, in a univariate setting, to AD and related intermediate traits such as quantitative measures of AD neuropathology and each individual’s trajectory of cognitive decline. Remarkably, m116, which is the module most strongly enriched for AD susceptibility genes, is positively associated with age but is not strongly associated with cognitive or pathologic measures of disease. Age is the strongest risk factor for AD, so it suggests that AD variants in m116 may primarily have a role in accelerating microglial aging and may not have strong, direct effects on β-amyloid and tau pathologies or cognitive decline beyond what is accounted for by advancing age.

**Table 2.** P-values for associations of modules to clinical and pathologic traits

| MODULE | ACTIVATED<br>MICROGLIA | PATHOLOGIC<br>AD | CLINICAL<br>AD | COGNITIVE<br>DECLINE | NEURITIC<br>PLAQUES | DIFFUSE<br>PLAQUES | NEUROFI-BRILLARY<br>TANGLES | TAU | B-AMYLOID | SEX | AGE |
| --- | --- | --- | --- | --- | --- | --- | --- | --- | --- | --- | --- |
| m116 | 0.042 | 0.16 | 0.15 | 0.2 | 0.091 | 0.61 | 0.68 | 0.58 | 0.016 | 0.013 | 0.0049 |
| m115 | 0.041 | 0.47 | 0.16 | 0.094 | 0.099 | 0.66 | 0.77 | 0.87 | 0.022 | 0.0044 | 0.4 |
| m114 | 0.02 | 0.0069 | 0.0045 | 0.0035 | 0.00031 | 0.055 | 0.24 | 0.024 | 0.00015 | 0.00016 | 0.5 |
| m113 | 0.63 | 0.43 | 0.026 | 0.085 | 0.1 | 0.28 | 0.75 | 0.87 | 0.0043 | 0.13 | 0.11 |
| m5 | 0.00042 | 0.005 | 0.068 | 0.014 | 0.0019 | 0.00036 | 0.43 | 0.0028 | 0.0064 | 0.00015 | 0.42 |

To replicate these associations, we used an existing dataset of frontal cortex data from subjects with AD and subjects without AD and imposed our module definitions on their data^4^; since our quantitative measures of AD neuropathology are not available in these subjects, we replicate the association of modules m5 and m114 with a pathologic diagnosis of AD. Both m5 (p=5.05×10^−5^) and m114 (p=1.13×10^−4^) replicate in this repurposed case/control dataset derived from brain bank samples. In a second level of replication and to address the issue of whether some of these associations may be detectable at the epigenetic level as well, we imposed our module definitions on H3K9Ac (a marker of active promoters) ChIPseq data available on 669 ROSMAP subjects, and we found that all of the previously observed associations are also found at this level, with the exception of m116 and age (**Supplementary Table 7**).

### Assessing the microglial sub-network within the neocortex

AD-related traits are correlated to one another (**Supplementary Figure 4**), as are the five modules, and this makes the interpretation of simple univariate analyses challenging, particularly in teasing apart associations with β-amyloid and tau pathologies. To resolve the most likely set of direct associations between modules and traits, we assessed the conditional dependence between the modules and traits by simultaneously considering all five modules and all pertinent traits to identify the subset of direct module-trait associations that will guide further work. We summarize this analysis in **Figures 2a and b**, which begins to distinguish how different modules of the innate immune system may be related to the different phases of AD.

**Figure 2.**
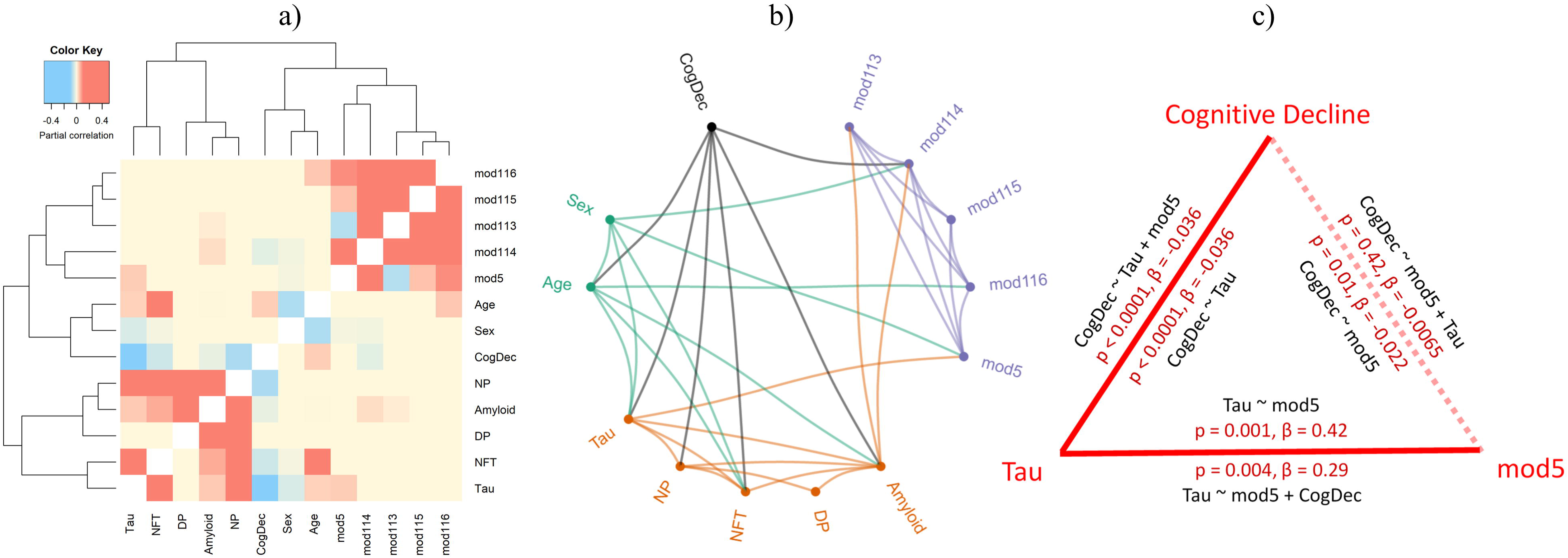
Relationships between gene modules and Alzheimer’s disease traits. The conditional dependencies between the expression of modules and Alzheimer’s disease traits are illustrated in a **(a)** heatmap and **(b)** network diagram. A link between a module and a trait is seen if there is evidence that they are associated after accounting for the behavior of all other modules and traits. **(c)** A diagram illustrating the relationships between cognitive decline, tau pathology and module 5 expression in isolation. For each edge in the triangle, the coefficient and its p-value are reported for the pair-wise and full models described.

There are multiple interesting observations in these results. First, we find female sex to be related to tau pathology; however, it is not associated with β-amyloid or cognitive decline in this multivariate model. Sex is also associated with two of the modules: m5 (p=0.00015) and m114 (p=0.00016), both of which are more highly expressed in women. Second, we report the expected association of worse cognitive decline as well as increased β-amyloid and tau burden with advancing age. Age is only associated with higher expression of m116 (p = 1.7×10^−30^). This result suggests that either the number or proportion of microglia in the neocortex (best captured by m116) is higher with age or that the expression of this set of genes is higher with age. The latter of these two hypotheses seems to be most likely as this m116:age association did not replicate in the H3K9Ac ChIPseq data, despite its strong effect size in the RNA data. So, while m116 is not directly related to the pathology measures or to cognitive decline, the overexpression of these genes in microglia of older individuals could contribute to the mechanisms behind aging, which is the single greatest risk factor for AD.

Looking at the other modules, we find that three are associated with pathologic measures; however, we begin to find distinct dissociations between β-amyloid (m113 and m114) and tau (m5) burden. m113 and m114 appear to be independently associated with β-amyloid pathology which typically accumulates early in the asymptomatic and early symptomatic phase of AD, and they are not directly associated with tau pathology. On the other hand, m5 is associated with the burden of tau pathology in an β-amyloid-independent manner (**Figure 2a and 2b**).

To begin to understand the magnitude of the effect of m5 and m114 on AD traits and to infer the most likely direction of the associations based on our cross-sectional data, we performed mediation analysis. Since cognitive decline, m114 and β-amyloid burden are all associated with one another, we tested for mediation, with the best fitting model finding that m114 is upstream of β-amyloid in the sequence of events as it influences cognitive decline through the accumulation of β-amyloid pathology (**Supplemental Figure 5c**). This is consistent with the observation that the asymptomatic accumulation of β-amyloid pathology occurs for many years before the presentation of symptoms, and our result suggest that one aspect of microglial function (m114) may therefore contribute to β-amyloid accumulation, consistent with functional studies of the *CD33* AD risk allele, which in turn contributes to cognitive decline.

In parallel, we evaluated m5 whose association with tau is very intriguing. When we add m5 to a model assessing the effect of sex on tau burden, we see that the effect of sex is diminished by 17%, suggesting that m5 may mediate part of the effect of sex on tau accumulation since sex determination occurs before the accumulation of late life pathologies. However, we highlight that m5 remains significantly associated with tau pathology after accounting for the effect of sex (p=1.58×10^−6^) (**Figures 2a**), so it must also be reflecting the effects of other mechanisms. To explore the possible sequence of events among m5 expression, cognitive decline and tau pathology, we performed a second mediation analysis which suggests that m5 expression precedes tau accumulation and its effect on cognitive decline may thus be mediated through the accumulation of tau pathology (**Figure 2c**). These results are consistent with previously described the putative role of microglia in a mouse model of tau pathology^20^.

### Role of m5 in the aging neocortex

Since m5 is distinct from m116 in the correlation structure of our RNAseq data but still very enriched for genes found in aged human microglia, we hypothesized that it may capture a subset of microglia with a particular function. We therefore turned to a phenotype that we had previously captured in our subjects^21^: the proportion of microglia with an activated stage III morphology based on immunohistochemical studies (**Figure 3a**). This neuropathologic measure is available in 104 subjects who also have RNAseq data. We observe a strong association between the expression of m5 and the proportion of microglia which are categorized morphologically as stage III, activated microglia (p-value = 0.00042, **Figure 3b, Supplementary Figure 6, Table 2**); in fact, out of all 47 cortical modules, m5 is the one most strongly associated with this trait. m5 is thus a transcriptional program related to microglial activation.

**Figure 3.**
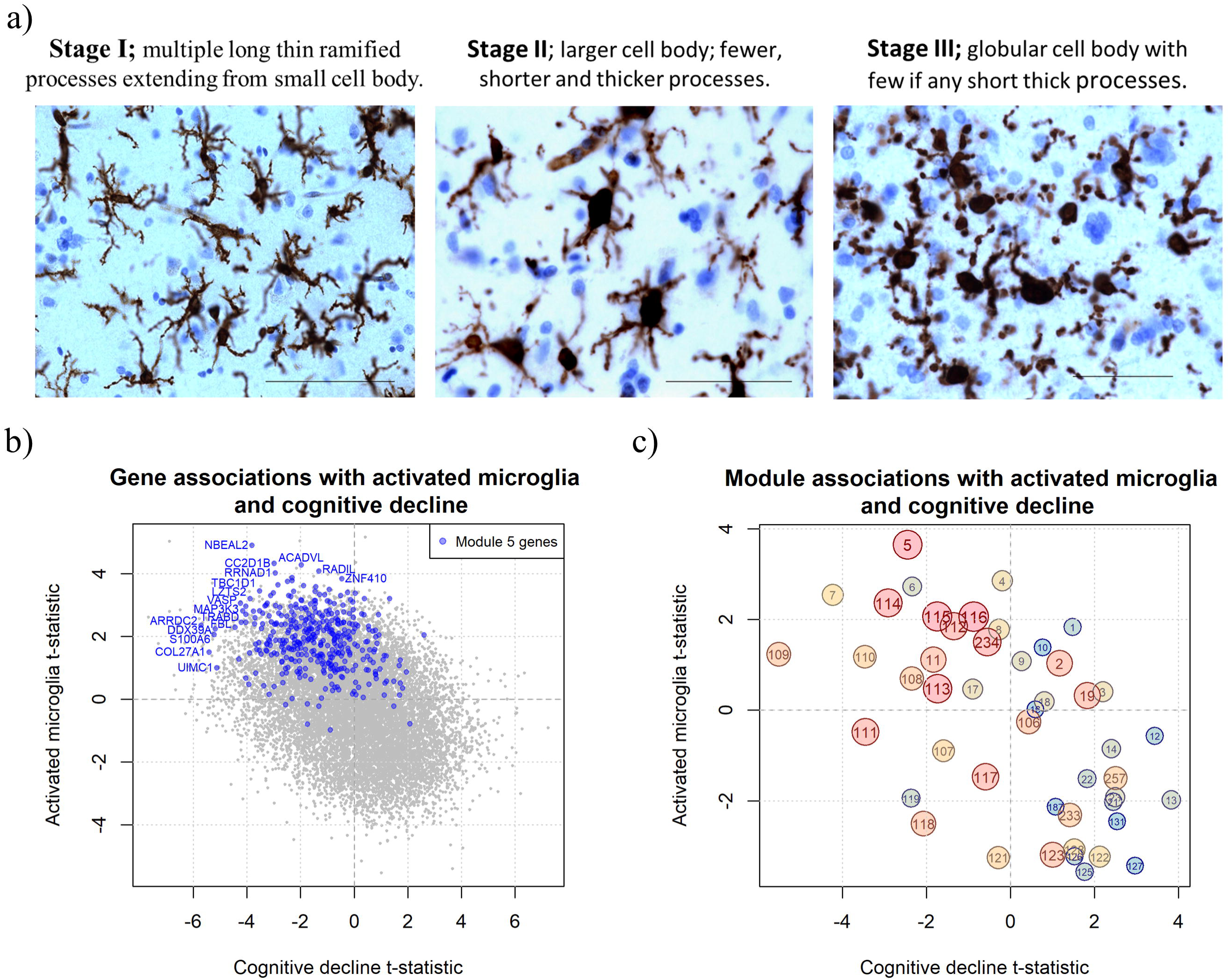
Association of genes and modules with microglia morphology. **(a)** Immunostaining showing stage I, II and III microglia morphologies. **(b)** The t-statistics from linear models which capture the associations between module expression with the proportion of Stage III microglia and a decline in cognition are compared. Modules are both colored (red to blue) and sized (big to small) by their enrichment for HuMiAged microglia signature genes. **(c)** The t-statistics from linear models which capture the associations of genes with activated microglia and a decline in cognition are compared. The genes in module 5 are highlighted in blue.

To take these analytic results to the next stage and confirm our results *in situ,* we selected a representative m5 gene, *VASP,* that statistically captured the effect of m5 on tau pathology and had antibodies available for immunofluoresence. At the single gene level, *VASP* RNA expression in cortical tissue is also strongly associated with activated microglial counts, tau burden and cognitive decline (**Table 3, Figure 3c**). Our modeling suggests that VASP should be expressed in microglia and should be expressed at higher level in microglia that have a morphology consistent with activation according to standard neuropathologic assessment (i.e. have a more globular, stage III morphology).

**Table 3.**
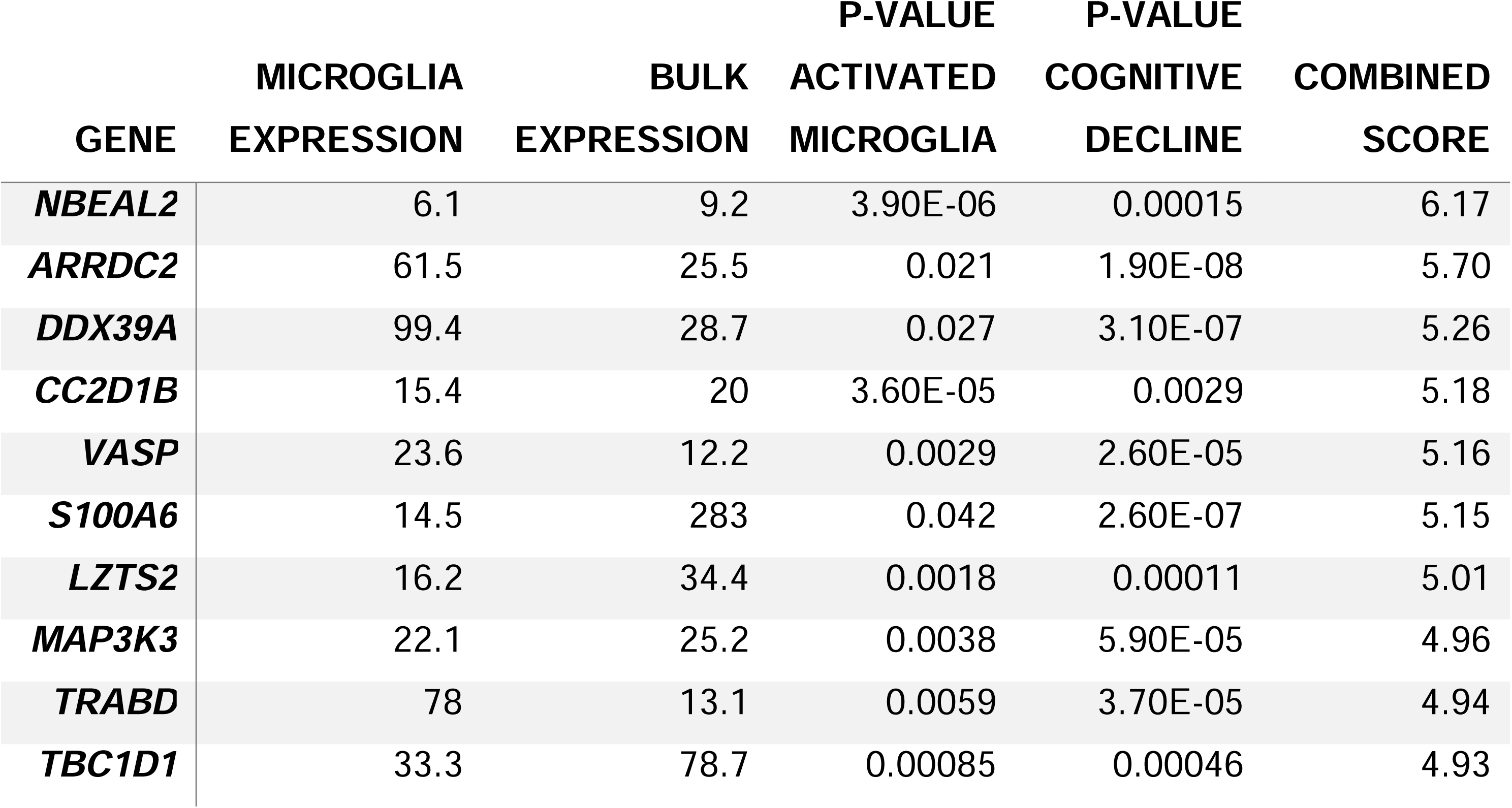
Top ten genes in module 5 that are most associated with activated microglia and cognitive decline.

| GENE | MICROGLIA | BULK | P-VALUE | P-VALUE | COMBINED<br>SCORE |
| --- | --- | --- | --- | --- | --- |
|  | EXPRESSION | EXPRESSION | ACTIVATED<br>MICROGLIA | COGNITIVE<br>DECLINE |  |
| *NBEAL2* | 6.1 | 9.2 | 3.90E-06 | 0.00015 | 6.17 |
| *ARRDC2* | 61.5 | 25.5 | 0.021 | 1.90E-08 | 5.70 |
| *DDX39A* | 99.4 | 28.7 | 0.027 | 3.10E-07 | 5.26 |
| *CC2D1B* | 15.4 | 20 | 3.60E-05 | 0.0029 | 5.18 |
| *VASP* | 23.6 | 12.2 | 0.0029 | 2.60E-05 | 5.16 |
| *S100A6* | 14.5 | 283 | 0.042 | 2.60E-07 | 5.15 |
| *LZTS2* | 16.2 | 34.4 | 0.0018 | 0.00011 | 5.01 |
| *MAP3K3* | 22.1 | 25.2 | 0.0038 | 5.90E-05 | 4.96 |
| *TRABD* | 78 | 13.1 | 0.0059 | 3.70E-05 | 4.94 |
| *TBC1D1* | 33.3 | 78.7 | 0.00085 | 0.00046 | 4.93 |

*VASP* has not been explicitly reported to be expressed in microglia previously, and so we first demonstrated that it is expressed in cells labeled with *TMEM119,* a protein proposed as a pan-microglial marker not expressed on infiltrating macrophages^22^ (**Figure 4a**). Interestingly, only a subset of *TMEM119+* microglia are *VASP+.* While *VASP* is reportedly expressed in human microglia^16^, it is also found to be expressed in fetal human astrocytes and to a much lesser extent in other cells of the CNS parenchyma. As shown in **Figure 4b**, we found *VASP* to be expressed in microglia in the vicinity of neuritic plaques, that consist of deposits of both β-amyloid and tau fibrils.

**Figure 4.**
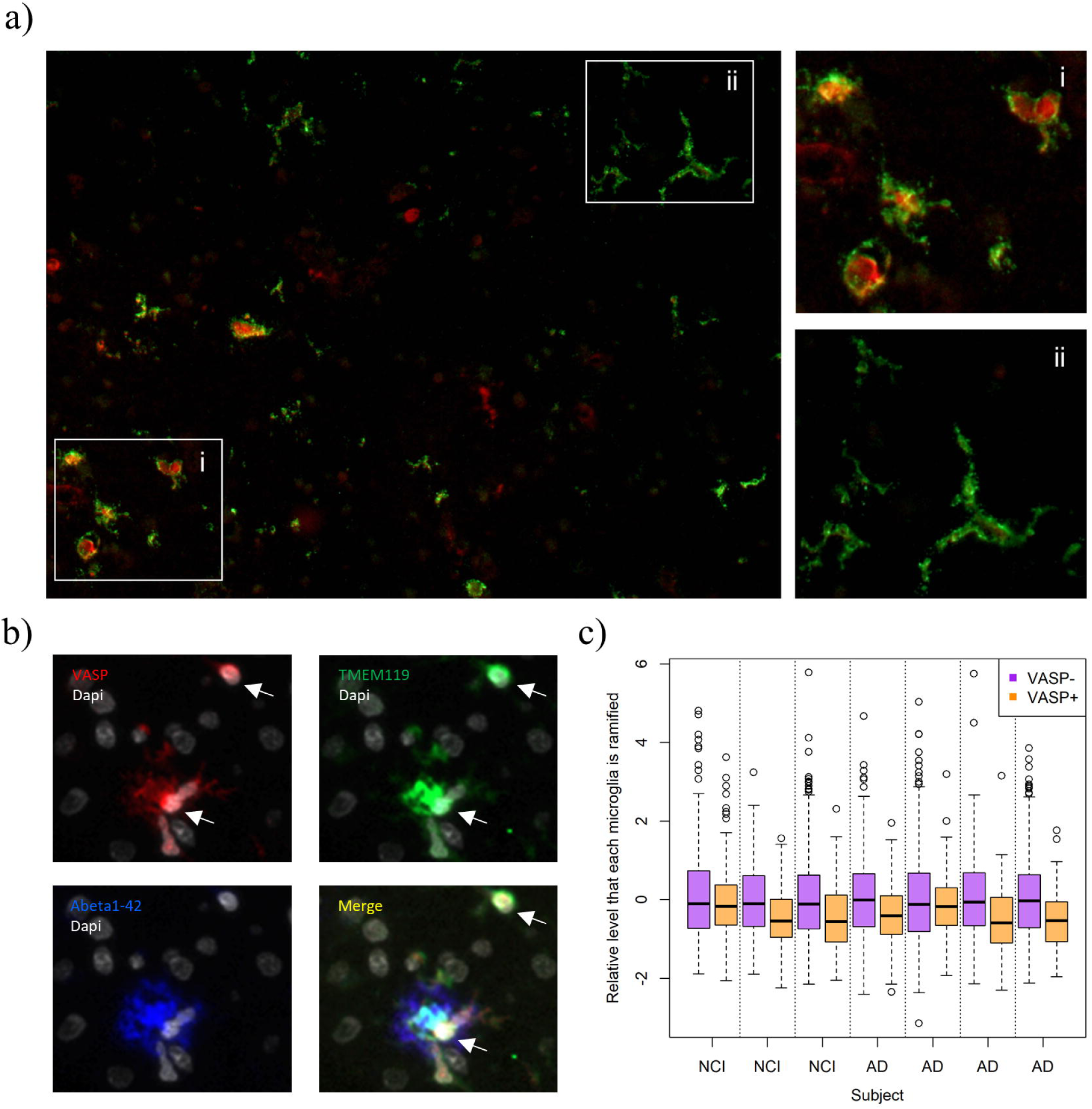
Immunofluorescence reveals VASP expression in TMEM119+ cells with higher expression in activated morphology. **(a)** Co-immunostaining of *VASP* (red) and *TMEM119* (green) in DLPFC. Nuclear staining of *VASP* in *TMEM119_+_* cells, with a less ramified morphology in *VASP_+_* (i) compared to *VASP-* (ii) cells. **(b)** Co-immunostaining of *VASP* (red), *TMEM119* (green), Aβ_1-42_ (blue) and *DAPI* (white). *TMEM119_+_* cells are close to the neuritic plaques expressed strongly *VASP.* (c) Quantification of ramified cell morphology for *VASP+* and *VASP-TMEM119+* cells.

To evaluate whether, as the m5 analyses suggest, *VASP* is a marker of activated microglia, we used automated image analysis ^23^ to assess whether the subset of *TMEM119+* cells that were also VASP+ had an activated morphology. To minimize bias, we used Cell Profiler to identify individual *TMEM119+* cells and then captured (1) the level of *VASP* expression by each cell and (2) several different morphologic features of each cell. Out of 4421 *TMEM119+* cells captured from 7 ROSMAP participants we find that *TMEM119+ VASP+* microglia are less ramified than *TMEM119+ VASP-* cells (p=2×10^−8^) (**Figure 4c**). The association between morphology appears to be similar among the limited number of pathologically defined AD subjects (n=4) and subjects with no cognitive impairment (n=3). Overall, these analyses confirm that m5, as estimated by its proxy marker *VASP,* is more highly expressed in morphologically activated microglial cells and that the *TMEM119+VASP+* cells are found in the vicinity of tau pathology.

## Discussion

We performed a detailed *in silico* dissection of five cortical transcriptional modules of human microglia in the aging human brain and discovered that they relate to different functional responses of microglia. In particular, we differentiated the role of m116 as a module reflecting microglial aging from those of m113 and m114 that are involved in promoting the accumulation of β-amyloid and of m5 that captures a morphologically-activated microglial state that contributes to the accumulation of tau. Thus, we begin to empirically differentiate groups of microglial genes that work together to accomplish specific, distinct functions. This regulatory architecture is present at the chromatin level: our H3K9Ac data both confirms the correlation of genes within RNAseq-defined modules and validates their association with AD-related traits.

Importantly, given our large sample size, we can identify different modules involved in β-amyloid and tau pathology. Our analyses suggest the existence of multiple microglial functional modules active in parallel within the same brain. In the case of m5, we pushed validation efforts further to confirm, *in situ,* the hypothesis that it reflects the presence of a subset of microglia that are more activated and found in the vicinity of neuritic plaques which contain tau fibrils in addition to amyloid. Diffuse plaques do not show this aggregation of activated microglia. Our data therefore implicate microglia in both β-amyloid and tau accumulation, and the three distinct transcriptional modules (m5, m113, and m114) are all present in the brain of aged individuals. This observation highlights the complexity that we face in developing immunomodulatory therapies for AD: they will have to be tuned to specifically engage or shut down a given transcriptional module while not exacerbating the ongoing perturbation of others. In the case of tau, we propose that the increased prevalence of activated microglia leads to more tau accumulation, suggesting that downregulation of this module may be a therapeutic option. The relatively specific enrichment of the JUN kinase pathway in m5 suggests that it may be regulated in a manner that is somewhat distinct from the other four modules and could provide a relatively specific target for drug development.

A surprising result is the lack of strong association between m116 and β-amyloid or tau: it is contrary to a simple narrative that this module, enriched for AD susceptibility genes from GWAS, exerts its effect primarily through increased accumulation of the neuropathologies that define AD. B-amyloid and tau burden only explain some of the variance of cognitive decline and AD dementia^9^, and our m116 results suggest that microglia may be involved in AD through mechanisms that remain elusive today. We clearly demonstrate the presence of an effect of aging on microglia, but this effect does not translate directly into pathology in our analysis. One thing to appreciate is that many AD GWAS often use samples of convenience as control subjects: these subjects are only coarsely characterized. Further, since AD incidence increases with advancing age, the majority of control subjects are likely to develop AD if they live long enough. Thus, the AD GWAS may have uncovered genes that have a strong effect on advancing the age of onset of AD and emerge as susceptibility genes because of the study design. Such an effect would tie in with our results where *TREM2, INPP5D,* and other genes in m116 are associated primarily with microglial aging.

One possibility is that the different transcriptional modules described here correspond to different microglia subpopulations. Given their sentinel and effector nature, the diversification of microglia phenotypes in specialized niches in the aged brain parenchyma burdened with β-amyloid and tau deposits is indeed a likely scenario. A longstanding observation is the morphological transformation of parenchymal microglia cells in and around the plaques in AD. Nonetheless, until now the molecular identity of the plaque associated (ameboid or stage III) activated microglia was obscure. This study takes the first steps to assign, though in an indirect way, a transcriptomic signature to a morphologically defined subpopulation of microglia. Future studies now in progress that utilize single cell RNA-sequencing approaches will allow us to understand how the transcriptional modules described here are represented in the population structure of microglia in the aged and AD human brain and what is the relationship of the subpopulations with clinical and histopathological features.

Nonetheless, our study has certain limitations; foremost amongst these is the fact that our pathologic measures are cross-sectional since they are obtained at autopsy. We therefore cannot comment on causality or on the exact sequence of events that is occurring in the human brain; mouse studies or *in vivo* microglial imaging will be necessary to help resolve such questions. Further, this neuroimmune network map should be seen as a first draft: microglia represent a minority of the cells found in the human cortex and have a relatively low quantity of mRNA in their cytoplasm. Thus, the level of expression of microglial genes, particularly those present only in a subset of microglia is likely to be underestimated or even absent from the tissue level data (e.g. CD33). Second generation maps derived from isolated microglia and single microglia transcriptional profiles will be essential to better understand the involvement of microglia in the pathophysiology of AD and the complex architecture of microglial subsets in the aged and AD brain.

Overall, we have generated an initial framework on which we and others can assemble additional data from *in vivo* and *in vitro* experimental models: when perturbing a particular gene of interest, we have to be cognizant of its membership to a given module and to the effect of such perturbations on different transcriptional programs found in microglia that may not be directly measured in an experimental system. The immune system is exquisitely modulated by a complex set of checks and balances in which AD genes such as *CD33, TREM1,* and *TREM2* are contributing to regulate the level of activation^24^, and, as seen in other immune functional programs, small differences in the level of receptor engagement or the presence of costimulatory molecules can result in dramatically different responses that can sometimes be the opposite of an anticipated response. By refining the set of microglial genes implicated in β-amyloid vs. tau pathology, we mark a big step forward in better targeting drug development programs both by proposing new targets and by defining novel outcome measures that can be used to assess the functional consequences of lead compounds.

## Acknowledgments

We thank the participants of ROS and MAP for their essential contributions and gift to these projects. This work has been supported by National Institute of Health (NIH) grants P330AG10161, U01 AG046152, R01AG16042, R01 AG036836, R01 AG015819, R01 AG017917 and R01 AG036547. The U01 AG046152 grant (to P.L.D.J. and D.A.B.) is a component of the AMP-AD Target Discovery Preclinical Validation Consortium, a program of the National Institute of Aging and the Foundation of the NIH.

## Author contributions

Study design: E.P, M.O, E.M.B, P.L.D.J. Sample collection: D.A.B. Data generation and quality control analyses: E.P., M.O., M.T, H-U.K, J.X., J.A.S, E.B, P.L.D.J. Analyses: E.P. Interpretation of results and critical review of the manuscript: E.P, M.O, M.T, H-U.K, J.X, C.C.W, D.F, C.G, L.B.C, S.M, J.A.S., D.A.B, E.M.B, P.L.D.J.

## Online methods

### ROSMAP cohort

The subjects profiled in this study are participants in one of two prospective cohort studies of aging, the Religious Orders Study (ROS)^6^ and the Memory and Aging Project (MAP)^7^ which are designed to be merged for joint analyses. These studies enroll non-demented individuals and include detailed, annual ante-mortem characterization of each subject’s cognitive status as well as prospective brain collection and a structured neuropathologic examination at the time of death. The study design of ROS and MAP yields an autopsy sample that includes a range of syndromic diagnoses and neuropathologic findings that are common in the older population. All brain autopsies, experiments and data analysis were done in compliance with protocols approved by the Partners Human Research Committee or the Rush University Institutional Review Board. The subjects in the study have an average age of 88, 61% meet criteria for pathologic AD by NIA Reagan criteria ^25^ and 64% are female.

### Description of RNA-Seq from ROSMAP

RNA was sequenced from the gray matter of dorsal lateral prefrontal cortex (DLPFC) of 542 samples, corresponding to 540 unique brains. These samples were extracted using Qiagen’s miRNeasey mini kit (cat. no. 217004) and the RNase free DNase Set (cat. no. 79254). RNA was quantified using Nanodrop. Quality of RNA was evaluated by the Agilent Bioanalyzer. All samples were chosen to pass two initial quality filters: RNA integrity (RIN) score >5 and quantity threshold of 5 ug (and were selected from a larger set of 724 samples). RNA-Seq library preparation was performed using the strand specific dUTP method ^26^ with poly-A selection^27^. Sequencing was performed on the Illumina HiSeq with 101bp paired-end reads and achieved coverage of 150M reads of the first 12 samples. These 12 samples will serve as a deep coverage reference and included 2 males and 2 females of non-impaired, mild cognitive impaired, and Alzheimer’s cases. The remaining samples were sequenced with a target coverage of 50M reads. The libraries were constructed and pooled according to the RIN scores such that similar RIN scores would be pooled together. Varying RIN scores results in a larger spread of insert sizes during library construction and leads to uneven coverage distribution throughout the pool.

The RNA-Seq data was processed by our parallelized pipeline. This pipeline includes trimming the beginning and ending bases from each read, identifying and trimming adapter sequences from reads, detecting and removing rRNA reads, and aligning reads to reference genome. The non-gapped aligner Bowtie was used to align reads to the transcriptome reference^28^, and RSEM was used to estimate expression levels for all transcripts^29^. The FPKM values were the outcome of our data RNA-Seq pipeline.

For normalization we first applied quantile normalization to the FPKM values and then used the combat algorithm^30^ to remove potential batch effects. Expression levels were quantified for 55,889 unique genes. GC and length bias effects were removed using a smoothed trimmed means of M-values. We placed a threshold for expression, only keeping 13153 genes with average FPKM greater than one.

### Description of Alzheimer’s disease related traits

### B-amyloid and Tau

To quantify β-amyloid and tau levels present in the brain at, tissue was dissected from eight regions of the brain: the hippocampus, entorhinal cortex, anterior cingulate cortex, midfrontal cortex, superior frontal cortex, inferior temporal cortex, angular gyrus, and calcarine cortex. 20μm sections from each region was stained with antibodies to the β-amyloid beta protein and the tau protein, and quantified with image analysis and stereology, as previously described^2,9,31,32^. Briefly, β-amyloid beta was labeled with an N-terminus-directed monoclonal antibody (10D5; Elan, Dublin, Ireland; 1:1,000). Immunohistochemistry was performed using diaminobenzidine as the reporter, with 2.5% nickel sulfate to enhance immunoreaction product contrast. Between 20 and 90 video images of stained sections were sampled and processed to determine the average percent area positive for β-amyloid beta. PHFtau was labeled with an antibody specific for phosphorylated tau (AT8; Innogenetics, San Ramon, CA; 1:1,000).

Between 120 and 700 grid interactions were sampled and processed, using the stereological mapping station, to determine the average density (per mm^2^) of PHFtau tangles. The scores across the eight regions were averaged, for β-amyloid and tau separately, to create a single summary measure for each protein. To create approximately normal distributions and facilitate statistical comparisons, we analyzed the square root of these two summary measures.

### Neuritic plaques and Neurofibrillary tangles

Neuritic plaque burden and Neurofibrillary tangle burden was determined by microscopic examination of silver-stained slides from 5 regions: midfrontal cortex, midtemporal cortex, inferior parietal cortex, entorhinal cortex, and hippocampus. The count of each region is scaled by dividing by the corresponding standard deviation. The 5 scaled regional measures are then averaged to obtain a summary measure for both neuritic plaque and Neurofibrillary tangle burden.

### Cognitive Decline

The ROS and MAP methods of assessing cognition have been extensively summarized in previous publications ^33-37^. Uniform structured clinical evaluations, including a comprehensive cognitive assessment, are administered annually to the ROS and MAP participants. Scores from 17 cognitive performance tests common in both studies were used to obtain a summary measure for global cognition as well as measures for five cognitive domains of episodic memory, visuospatial ability, perceptual speed, semantic memory, and working memory. The summary measure for global cognition is calculated by averaging the standardized scores of the 17 tests, and the summary measure for each domain is calculated similarly by averaging the standardized scores of the tests specific to that domain. To obtain a measurement of cognitive decline, the annual global cognitive scores are modeled longitudinally with a mixed effects model, adjusting for age, sex and education, providing person specific random slopes of decline. The random slope of each subject captures the individual rate of cognitive decline after adjusting for age, sex, and education. Further details of the statistical methodology have been previously described^38^.

### Microglia morphology

Immunohistochemistry for microglia was performed using an Automated Leica Bond immunostainer (Leica Microsystems Inc., Bannockborn IL) and anti-human HLA-DP, DQ, DR antibodies (clone CR3/43; DakoCytomation, Carpinteria CA; 1:100) using standard Bond epitope retrieval and detection. An investigator blinded to the clinical and pathologic data, outlined the cortical gray matter region of interest on each slide using a Microbrightfield Stereology System. The Stereo Investigator 8.0 software program was used to place a 1000 χ 750 μm sampling grid over the region and the program was engaged to sample 4.0% of the region with a 200 χ 150 μm counting frame at 400x magnification at interval grid intersection points. Using separate tags for stage 1, 2 and 3 microglia, the operator marked the microglia at each intersection point. These counts were then upweighted by the stereology software to estimate total number of microglia (stage 1, 2 and 3) in the defined area. Different stages of activation from least (stage 1) to most (stage 3) activated can be defined morphologically; when these cells become activated, their long fine processes contract and thicken and the cell body adopts a larger more rounded cellular conformation.

### Definition of Microglia genes

An aged human microglia signature (HuMiAged gene set) of 1081 microglia related genes were defined by comparing the expression of genes in DLPFC isolated microglia to their expression level in the bulk DLPFC tissue. The identified genes had four-fold larger expression in the microglia samples relative to the tissue level samples and average FPKM greater than one in the microglia samples. These microglia related genes can be viewed in the microglia vs bulk space in Figure 1.

### Definition of modules from ROSMAP

The SpeakEasy algorithm^39^ was used to derive gene modules from normalized gene expression data. Consensus clustering results from 100 initializations of the SE algorithm yielded 257 modules, 47 of which contained at least 20 gene members (meaning modules are assigned for 98% of genes) and were examined in downstream analyses. Pseudo-expression values for each module were calculated by taking the mean expression level of all genes assigned to that module after the expression data has been standardized for each gene.

### Modules enriched for microglia genes

A series of hypergeometric tests were used to test whether modules contained more microglia related genes than expected by chance. All 13153 genes that had FPKM greater than 1 in the tissue level samples were used as the background universe for the tests.

### Pathway analysis of modules

A series of hypergeometric tests were used to test whether modules contained more genes in any of the Gene Ontology biological process processes ^40^ than expected by chance. Only biological processes with more than 20 annotated genes or less than 500 genes were tested. Genes that were both in at least one biological process and module were used as the background universe for the tests.

### GWAS hits

#### IGAP enrichment

A series of hypergeometric tests were used to test whether modules contained more genes with at least one probe that was 50KB up or down-stream of the gene that was one nominally significant probe in IGAP ^18^ than expected by chance. Genes that were in the module definition and contained at least one IGAP probe were used as the background universe for the tests.

### Associations with Alzheimer’s related traits

#### Testing association with traits

Simple linear regression analysis was used to associate either gene expression or average module expression to several variables that measured the neuro-pathology in the AD brain and cognitive decline. These included either the numbers of neuritic plaques (NP), neurofibrillary tangles (NFT), the amount of β-amyloid or tau and the rate of cognitive decline (cogDec). All the associations were adjusted for age, sex, study (ROS or MAP), RNA Integrity number (RIN) and post-mortem interval (PMI).

#### Replication of association in Zhang et al.

We replicated the association of the immune modules with Alzheimer’s disease pathology in an independent microarray gene expression data from cortex (DLPFC) from a previous study by Zhang and colleagues^41^. Data was downloaded from GEO with accession number GSE44772. The expression data was quantile normalized and corrected for age, gender, post-mortem interval, pH, RIN and batch. Simple linear regression was then used to test for association with diagnosis of Alzheimer’s disease.

#### Partial correlation analysis

Partial correlations were used to further disentangle the highly correlated modules and traits. A partial correlation between a trait and a module is the correlation between the trait and module after accounting of the behavior of all other traits and modules. Graphical lasso ^42^ was used to set small partial correlations to zero and was tuned using repeated 10-fold cross-validation to estimate the penalty parameter.

### Activated microglia validation

Six μm sections of formalin-fixed paraffin-embedded tissue from the DLPFC were used to stain *TMEM119* (Sigma Aldrich), *VASP* (Santa Cruz Biotech) and β-amyloid 1-42 (Biolegend). Immunohistochemistry was performed using citrate as antigen retrieval. The sections are blocked with blocking medium (8% of horse serum and 3% of BSA) and incubated overnight at 4°C with primary antibodies. Sections are washed with PBS and incubated with fluorochrome conjugated secondary antibodies (Thermo Fisher) and coverslipped with anti-fading reagent containing Dapi (P36931, Life technology). Photomicrographs are captured at X20 magnification using Zeiss Axio Observer.Z1 fluorescence microscope and exported to Image J imaging software (NIH, Maryland, USA). The images are analyzed in collaboration with the Broad Institute’s Imaging platform, using CellProfiler and CellProfiler Analyst systems to measure and classify cells according to the cell morphology. *TMEM119_+_* cells and *VASP_+_* are first identified using the default thresholding, overlaps of these cells are then considered to identify *TMEM119+ VASP-* cells and *TMEM119_+_ VASP_+_* cells. The compactness parameter, the variance of the radial distance of the object’s pixels from the centroid divided by the area, from CellProfiler is then compared between *TMEM119_+_ VASP-* cells and *TMEM119_+_ VASP_+_* cells. A random effects model accounting for between subject variability is used to assess the significance of differing compactness between the *TMEM119_+_ VASP-* cells and *TMEM119_+_ VASP_+_* cells.

